# An Eligibility-Aware Pipeline for Robust ITS Diagnostics in Fungi: A Cacao Case Study with Generalizable Rules

**DOI:** 10.1101/2025.11.01.686068

**Authors:** Seunghyun Lim, Insuck Baek, Jishnu Bhatt, Rakesh K. Upadhyay, Sookyung Oh, Minhyeok Cha, Moon S. Kim, Lyndel W. Meinhardt, Ezekiel Ahn

## Abstract

Accurate fungal ITS diagnostics rely on public sequence archives, but heterogeneous record lengths, especially frequent truncation before the LSU/28S segment, cause naive *in silico* benchmarking to conflate primer performance with database incompleteness. To resolve this persistent “denominator error,” we present an open and fully reproducible “eligibility-aware” framework. Our pipeline first establishes eligibility by confirming both primer sites are present before applying bench-realistic performance rules, including a strict penalty for 3’-terminal mismatches. It further provides mechanistic insights by analyzing binding-site conservation and uses a rarefaction-based approach to guide efficient quality control as databases grow. We demonstrate the framework’s utility using the cacao pathosystem, a context where rapid differentiation of the fungal pathogen *Moniliophthora* from symptomatically similar oomycetes is critical. The result is a robust, field-ready diagnostic decision tree operable under a single touchdown (TD) PCR program. By providing a transparent and barcode-agnostic template, our eligibility-aware approach offers a significant methodological advance for designing and validating molecular assays in mycology and beyond.

## 1 Introduction

The nuclear ribosomal internal transcribed spacer (ITS) remains the de facto barcode for fungi. Yet, the public archives that underpin primer design and diagnostic benchmarking are heterogeneous in curation and read length. In particular, many records end within ITS2 and do not extend into the LSU/28S segment where canonical reverse primers (e.g., ITS4) bind. As a result, naive in silico PCR, evaluated against all accessions, confounds primer performance with database incompleteness, creating a persistent denominator error (Nilsson et al., 2008; Schoch et al., 2012; Taylor et al., 2016).

We address this by formalizing an eligibility-aware accounting that separates (i) eligibility, whether both primer sites are present and oriented to yield a plausible product, from (ii) performance—bench-realistic success under strict rules (≤ 2 total mismatches across the pair and no 3′-terminal mismatch) within a defined size window. We complement these rules with binding-site information content/entropy to mechanistically link 3′-end conservation to amplification outcomes, and with rarefaction over primer-binding “haplotypes” to set efficient sample sizes for routine quality control (Bokulich & Mills, 2013; Gardes & Bruns, 1993; Ihrmark et al., 2012; Toju et al., 2012; White, 1990).

To illustrate practical use in a plant-health context, we structure a worked example in cacao (*Theobroma cacao*) where rapid identification of a basidiomycete target (*Moniliophthora* spp.) (Bailey et al., 2018; Meinhardt et al., 2008) is complicated by symptomatically similar agents (Ali et al., 2016; Franco et al., 2019; Holmes et al., 2003). The example is didactic: the framework itself is fungal-first and barcode-agnostic, not tied to properties unique to cacao.

Here, we present an open, eligibility-aware software framework that implements these principles under a conservative, bench- and realistic rule set. The pipeline first establishes eligibility by confirming that both primer sites are detectable (IUPAC-aware) and oriented to produce a positive-length product. It then scores performance based on strict criteria: no 3′-terminal mismatch on either primer, ≤ 2 total mismatches across the pair, and amplicon length is constrained within a realistic window (Taylor et al., 2016; White, 1990). From accession-level FASTA inputs and a primer catalog, the tool produces: (1) accession-resolved calls with explicit reason codes (strict hit, fail_MMs, fail_3′, ineligible class); (2) binding-site sequence logos and positional Shannon entropy to mechanistically link performance to 3′-end conservation (Bokulich & Mills, 2013; Toju et al., 2012); (3) rarefaction of binding-site haplotypes to set efficient sample sizes for ongoing QC; and (4) a lineage-aware decision tree derived from these outputs; for wet-lab deployment we recommend a single touchdown PCR program (TD 62→45 °C) to mitigate mis-priming and absorb modest ΔTm (Don et al., 1991; Rose et al., 1998).

Although our worked example involves separating *Moniliophthora* (fungi) from *Phytophthora* (oomycetes, which produce symptomatically similar black-pod lesions) in cacao with a minimal ITS panel, the logic is barcode-agnostic. It generalizes to other primer-delimited markers relevant to plant systems (e.g., rbcL for the ribulose-1,5-bisphosphate carboxylase/oxygenase large subunit gene and matK for the maturase K gene in plants, and COI for the cytochrome c oxidase I gene, where appropriate) and to pathosystems subject to truncation-heavy public archives (Nilsson et al., 2008; Schoch et al., 2012). By enforcing denominator discipline and publishing accession-level audit trails, the framework prevents routine interpretive errors, clarifies why recommended assays work (via 3′-end conservation), and renders routine QC tractable through principled down-sampling. In doing so, it supports plant disease surveillance and crop biosecurity while retaining the portability and transparency of PCR-based workflows (Franco et al., 2019). As a community resource, we release all code and accession-resolved outputs. We recommend reporting eligibility-aware and naive coverages side by side with explicit ineligibility reasons. This minimum reporting standard prevents misinterpretation and promotes robust, reusable ITS diagnostics across fungal systems.

## 2 Materials and Methods

### 2.1 Pipeline Architecture

The software is organized as a linear pipeline of five composable modules that run sequentially with sensible defaults. The workflow proceeds from an initial Eligibility Scan to the application of Strict Calls, followed by the generation of Logos and Entropy plots, a Primer-Site Rarefaction analysis, and finally the construction of a Decision-Tree Summary. Each step produces auditable outputs (CSV files), supporting transparent review and adaptation to other pathosystems.

### 2.2 Implementation and Reproducibility

The pipeline is implemented in Python 3 (CPython 3.10–3.11), and all dependencies are pinned in requirements.txt to support reproducible installations. To ensure full reproducibility, we provide two primary routes:

1. A Jupyter notebook (notebooks/run_all.ipynb) that reproduces the analyses from the raw input data.
2. A command-line interface (scripts/scan_pairs.py) for batch processing of large FASTA files (Peng, 2011).

All randomization procedures use a fixed seed (--seed 17) to ensure results are bit-for-bit reproducible.

### 2.3 Core Modules and Functionality

The framework consists of five primary modules accessible via both the command-line and a Python API:

i. Eligibility Scan: This IUPAC-aware site-finder takes FASTA files and a primer catalog as input (Table S1). It locates the optimal forward and reverse primer binding sites within each accession, accommodates degenerate symbols, and verifies a valid product orientation (Rice et al., 2000; Walters et al., 2011). It produces auditable outputs, including detailed results for each accession and summary tallies by species.
ii. Strict Calls: Using the output from the eligibility scan, this module applies a set of strict, bench-realistic scoring rules to all eligible sequences (default: ≤ 2 total mismatches, no 3′-terminal mismatch, and within a defined amplicon window) (Kwok et al., 1990; Stadhouders et al., 2010).
iii. Logos and Entropy: To visualize binding-site conservation, this module generates sequence logos and plots of per-position Shannon entropy for the primer binding words. The plots are annotated to mark the critical 3′-terminal base, providing a mechanistic link between sequence conservation and the no-3′-mismatch rule (Schneider & Stephens, 1990; Tareen & Kinney, 2020).
iv. Primer-Site Rarefaction: This module provides a tool for scalable quality control. It treats each unique primer binding sequence as a “haplotype” and computes a rarefaction curve to estimate N_95_, the sample size required to capture ~95% of the observed haplotype diversity (Chao & Jost, 2012; Gotelli & Colwell, 2001).
v. Decision-Tree Summary: The pipeline produces the summary data tables described above. These outputs were then used to derive a compact, lineage-aware PCR decision tree that recommends a default short amplicon for each target lineage. The decision-making logic derived from these outputs is presented visually in the form of flowcharts. For laboratory deployment, we recommend a single touchdown PCR program to mitigate mis-priming and absorb modest ΔTm (Don et al., 1991).

### 2.4 Work Example and Sequence Data

Our worked example used public ITS accessions for the fungal pathogens *M. perniciosa* (n = 182) and *M. roreri* (n = 186). As a diagnostic comparator, we also included the oomycete pathogens *P. palmivora* (n = 1,221) and *P. megakarya* (n = 82), which cause similar black pod symptoms in cacao. All FASTA records were sourced from GenBank, normalized to uppercase, and analyzed without prior trimming to preserve the signal of archive truncation.

## 3 Results

### 3.1 Exhaustive *in silico* Screening Produces a Minimal, Lineage-Aware Panel

We evaluated all biologically sensible ITS forward–reverse combinations for the four focal taxa under our default strict rule set, as defined in the Methods. To make the denominator effect explicit, we report for each species × pair the naive coverage (strict hits as a percentage of all accessions), the eligibility-aware coverage (strict hits among eligible accessions), and their difference (“Δ-coverage, pp”, Table S2). Sensitivity to scoring thresholds is summarized in Supplementary Data 1. Coverage is reported among eligible accessions, those in which both primer sites are found with ≤ 4 mismatches and a valid product orientation/size. We also identify and exclude ineligible rows to prevent archive truncation from being misinterpreted as a primer failure. To prevent inflated denominators, counts for species-by-pair bars were computed after deduplication to one row per accession × pair. Unless otherwise noted, results are reported for fungi of interest; comparator lineages are included only to provide differential-diagnosis context and to expose archive-driven ineligibility patterns.

Figure 1 shows species-by-pair coverage (strict hits, eligible-not-strict, ineligible) for the 12 pairs most relevant to the recommended panel; the complete per-pair matrix remains in Supplementary Data 1. Figure 2 visualizes primer-site conservation with information-content logos (y-axis in bits; max = 2) and positional Shannon entropy, with the 3′ base marked by a dashed line. Figure 3 presents primer-site rarefaction. Figure 4 consolidates these outputs into the decision-tree panel. All row-level calls and species-level tallies (for every pair tested) are provided in Supplementary Data 1.

**FIGURE 1.**
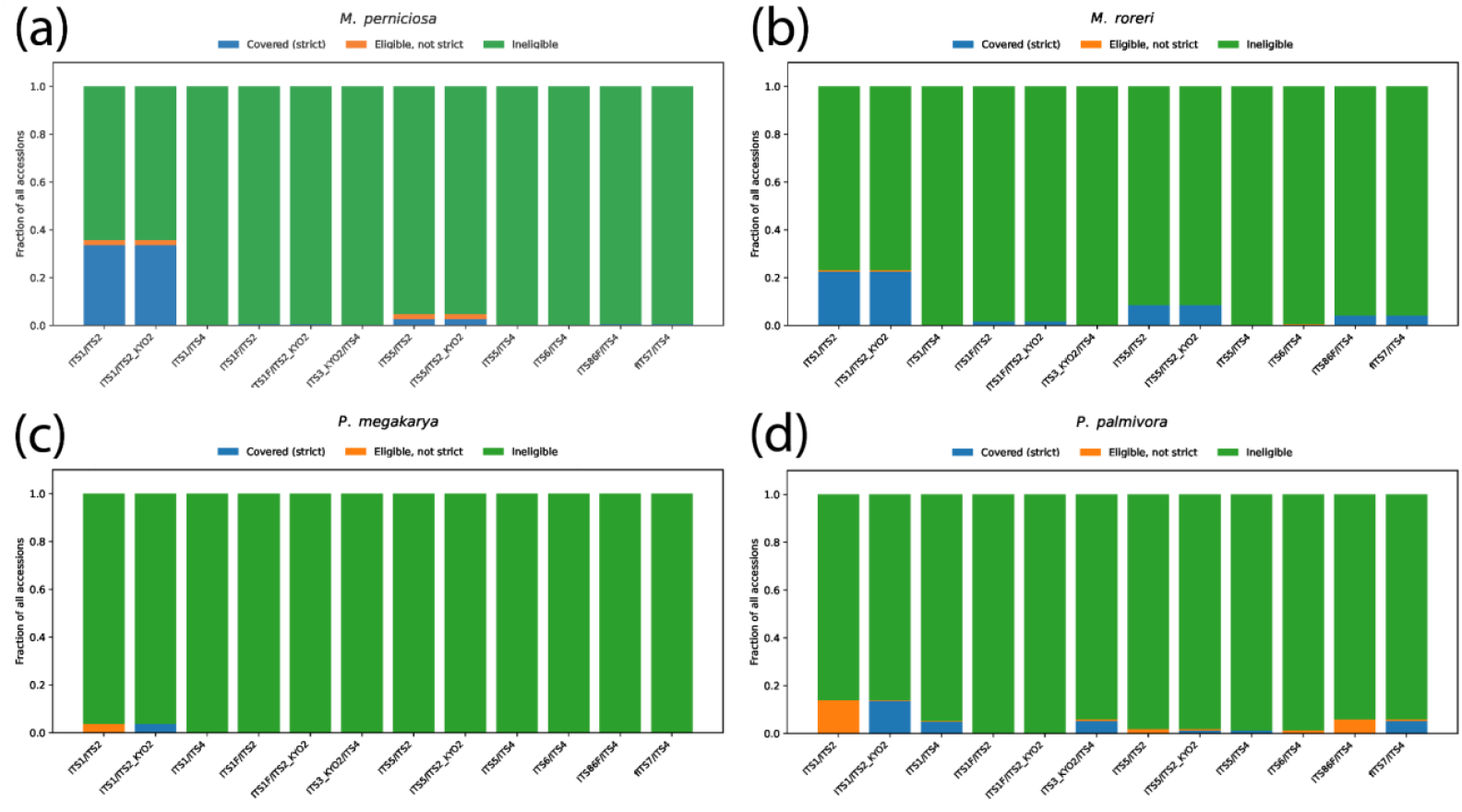
Coverage across primer pairs by species after deduplication to one row per accession × pair. (a) *M. perniciosa*, (b) *M. roreri*, (c) *P. megakarya*, (d) *P. palmivora*. Stacked bars show the fraction of all accessions: strict hits (blue), eligible but not strict (orange), and ineligible (green).

**FIGURE 2.**
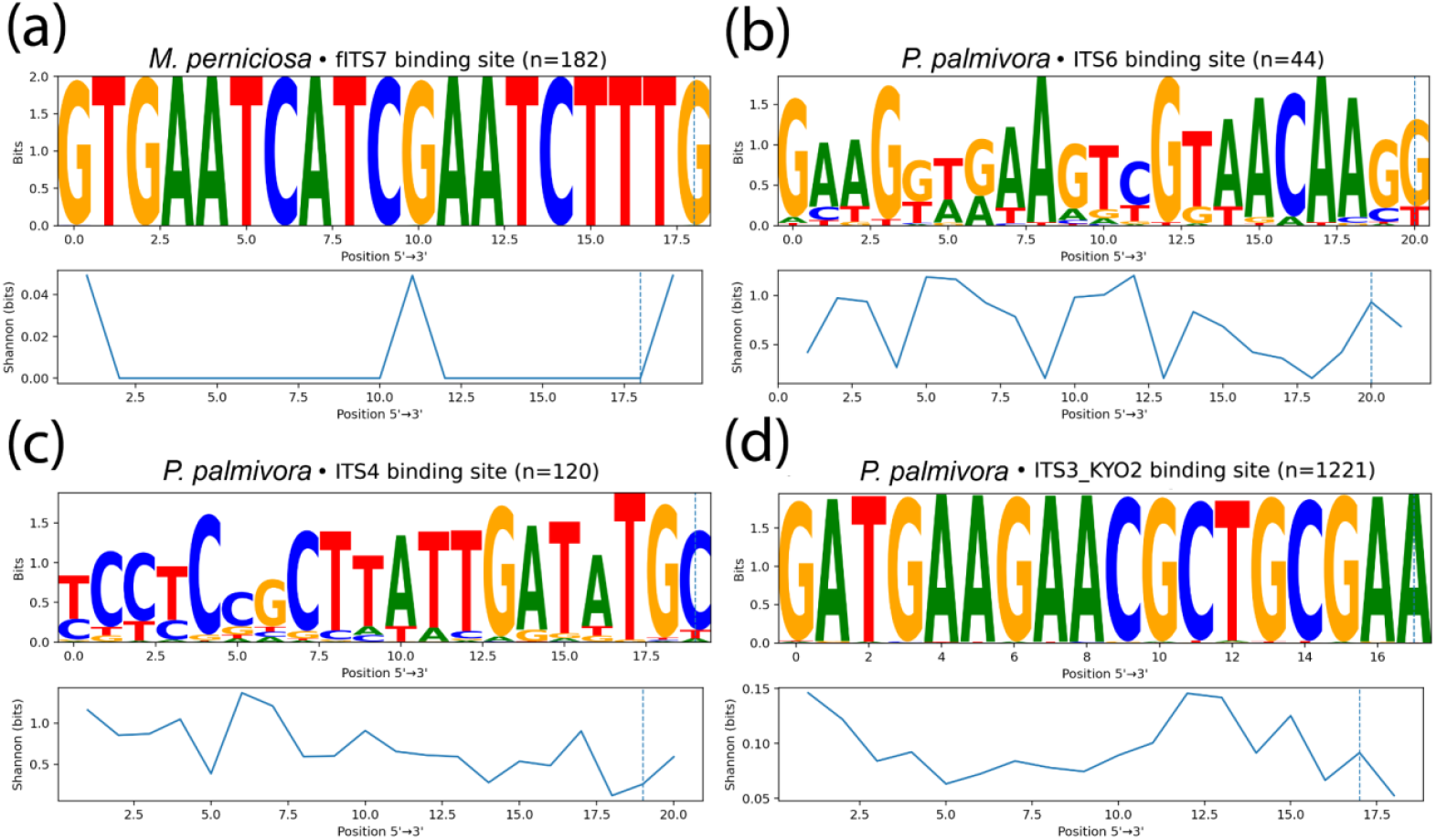
Primer binding-site logos (information content; bits) and positional Shannon entropy (5′→3′). (a) *M. perniciosa* fITS7 forward (n = 182), (b) *P. palmivora* ITS6 forward (n = 44), (c) *P. palmivora* ITS4 reverse (rendered on the forward strand; n = 120), (d) *P. palmivora* ITS3_KYO2 forward (n = 1,221). The dashed line marks the primer’s 3′ terminal base; low entropy/high information content at this position is consistent with high strict-hit rates under the no-3′-mismatch rule.

**FIGURE 3.**
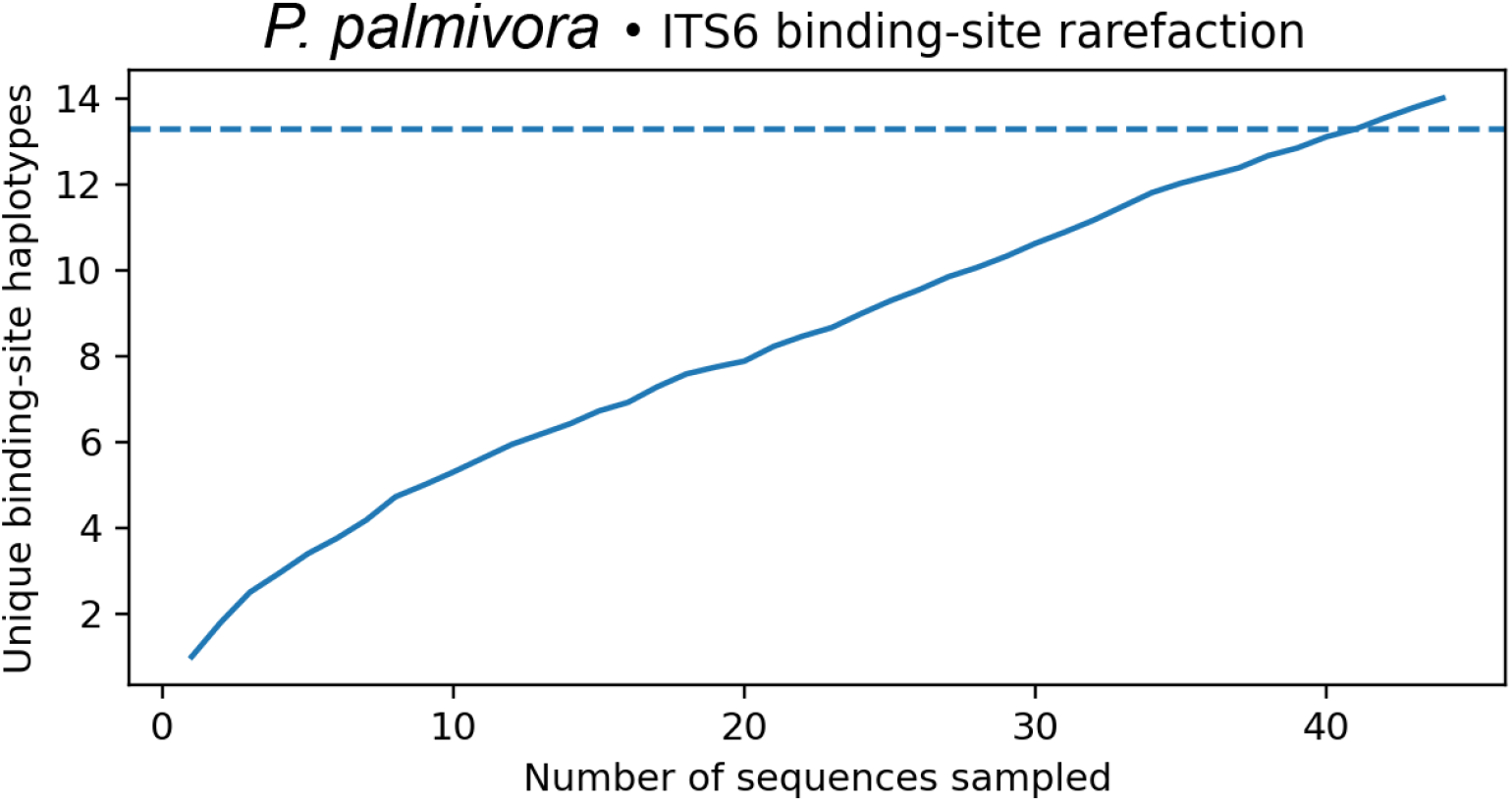
Rarefaction of *P. palmivora* ITS6 binding-site haplotypes. Mean cumulative unique count across 50 random permutations (solid line). The dashed line marks 0.95 × the total unique haplotypes observed; the curve intersects at ~ 41/44 sequences.

**FIGURE 4.**
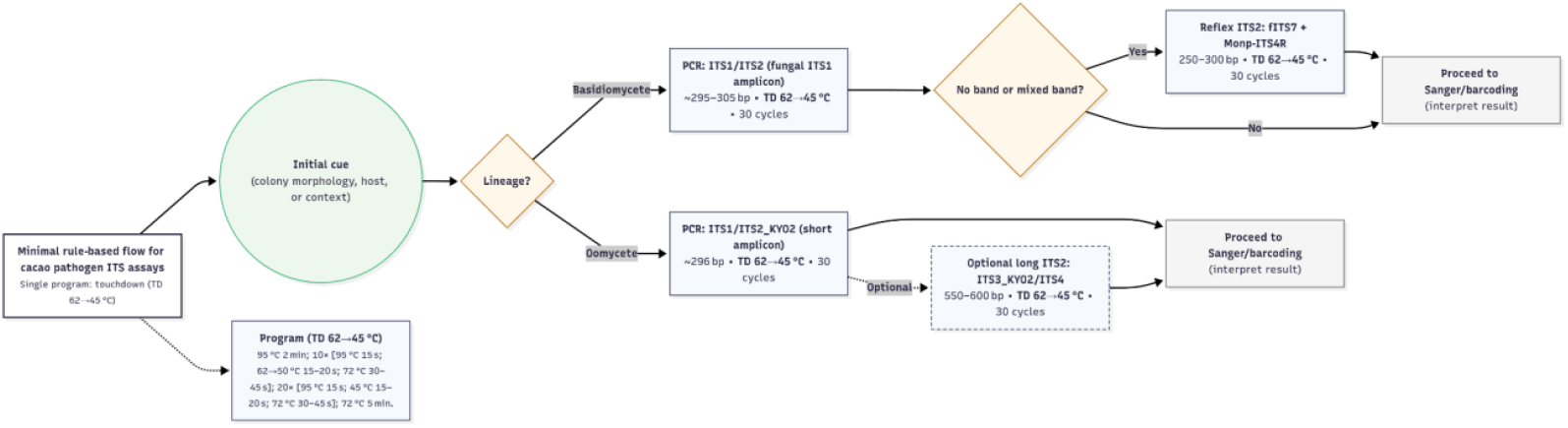
Minimal, rule-based decision tree for cacao pathogen ITS diagnostics under a single TD program. Top branch: *Moniliophthora* → ITS1/ITS2; reflex to fITS7 + Monp-ITS4R when LSU allows. Bottom branch: *Phytophthora* → ITS1/ITS2_KYO2; optional long ITS2 readout ITS3_KYO2/ITS4 when LSU allows. Program A/B fixed-anneal alternatives appear in Figure S2.

To quantify the denominator error, we calculated Δ-coverage values (Table S2). A paired summary shows a large positive Δ for assays whose reverse site is often missing (e.g., *M. roreri* fITS7 (gITS7 variant)/ITS4: Δ = 96 pp; *M. perniciosa* fITS7/ITS4: Δ = 99 pp), and likewise for *P. palmivora* ITS1/ITS2_KYO2, where many records lack the ITS1 forward block (Δ = 85 pp). By contrast, Δ is modest for ITS6/ITS4 (Δ = 12 pp), reflecting both a small eligible pool and lower strict-hit rates within that pool (Table S2).

### 3.2 Basidiomycetes (*Moniliophthora*): ITS1/ITS2 is a Robust Primary Assay; Long ITS2 Readouts Depend on LSU

For *M. perniciosa*, ITS1/ITS2 covered 61 of 65 eligible accessions (93.8%) under strict rules, with 117 additional rows being ineligible (primarily due to the absence of forward sites for this pair). In contrast, fITS7/ITS4 (an ITS2 forward with an LSU-anchored reverse) had a single eligible record that was a strict hit (1/1) and 181/182 ineligible rows due to missing ITS4 sites— i.e., lack of the LSU flank, not primer failure. *M. roreri* showed the same pattern (ITS1/ITS2: 42/43 strict; fITS7/ITS4: 8/8 strict among eligibles, but most rows were ineligible for the long assay for the same reason). These results motivate a short ITS1 amplicon (ITS1/ITS2) as the default for *Moniliophthora*, reserving an ITS2 readout (e.g., fITS7 + a suitable LSU reverse) for accessions verified to extend into 28S (Figure 1a–b; Supplementary Data 1). As shown in Figure 1, the large ineligible fractions for these long ITS2-forward + ITS4 assays are primarily driven by the absence of the LSU-anchored ITS4 site in public archives, rather than by primer failure itself. Indeed, fITS7/ITS4 showed Δ ≈ 99 pp in *M. perniciosa* and 96 pp in *M. roreri* because almost all records lacked the LSU-anchored ITS4 site (Table S2).

### 3.3 Oomycetes (*Phytophthora*): Short ITS1 Assays Generalize; Long ITS2 Readouts are Useful When LSU is Present

In *P. palmivora*, ITS1/ITS2_KYO2 covered 167/169 eligible (98.8%; 2 fail_MM; 0 fail_3′), with 1,052 additional rows ineligible (driven by archive structure relative to the reverse primer for other pairs). The canonical oomycete baseline ITS6/ITS4 had 17 eligible and 2/17 strict hits (11.8%), with 1,204 ineligible rows. The longer assay ITS3_KYO2/ITS4 expanded the eligible pool to 73 and covered 64/73 (87.7%) (8 fail_MM; 2 fail_3′), with 1,148 ineligible. These patterns recommend ITS1/ITS2_KYO2 as the default short-amplicon assay for *Phytophthora*, with ITS3_KYO2/ITS4 as an optional longer readout when LSU is present. *P. megakarya* was sparsely represented (short assay 3/3 strict; long assays had no eligible accessions in the present dataset) (Figure 1c–d and Table 1). For *P. palmivora* ITS1/ITS2_KYO2, Δ was ~85 percentage points, whereas the long ITS3_KYO2/ITS4 assay showed a smaller Δ (~83 pp) because its eligible pool was larger; ITS6/ITS4 had a very small eligible pool and thus a large Δ despite few strict hits overall (Table S2).

**TABLE 1.**
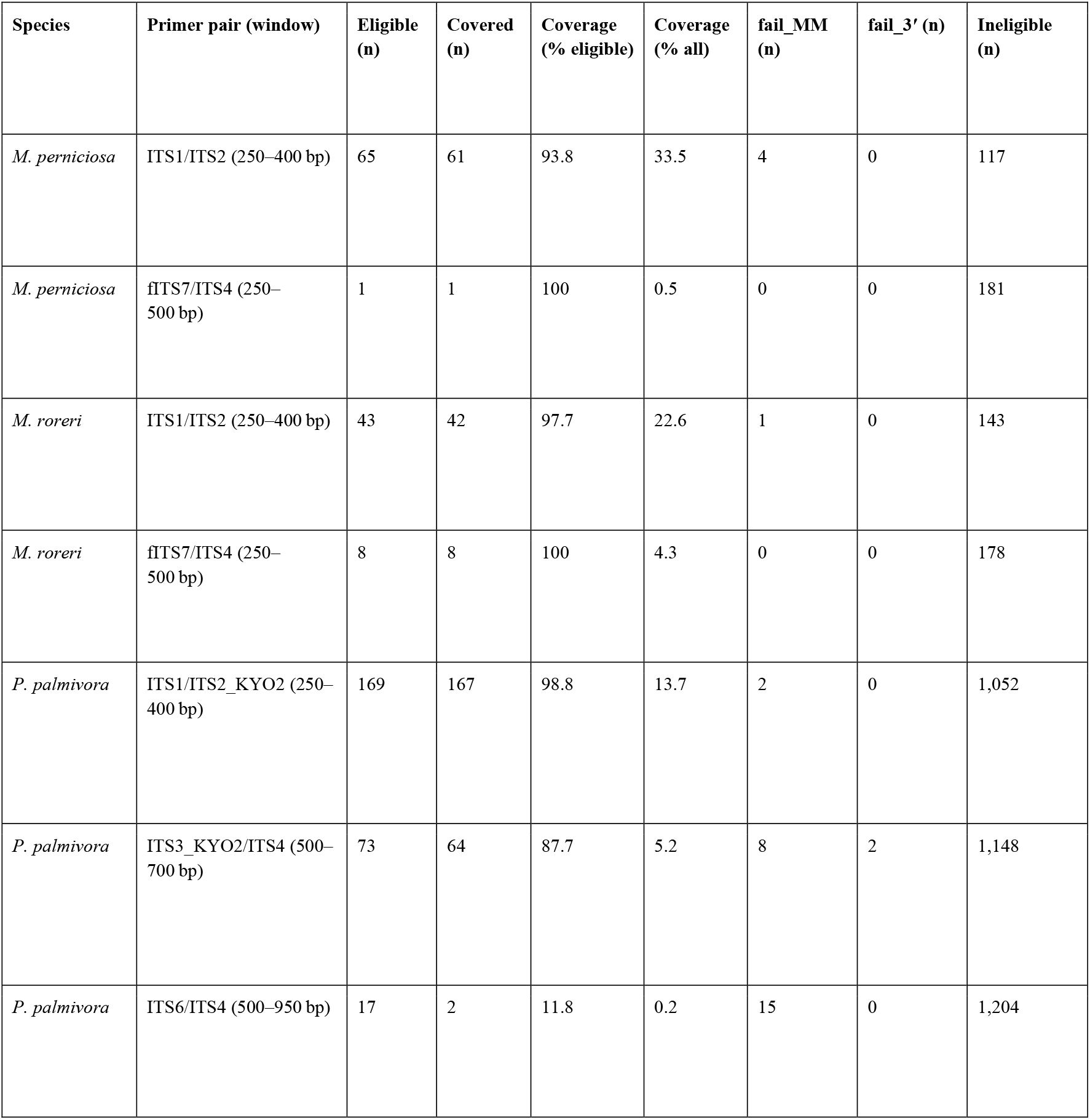
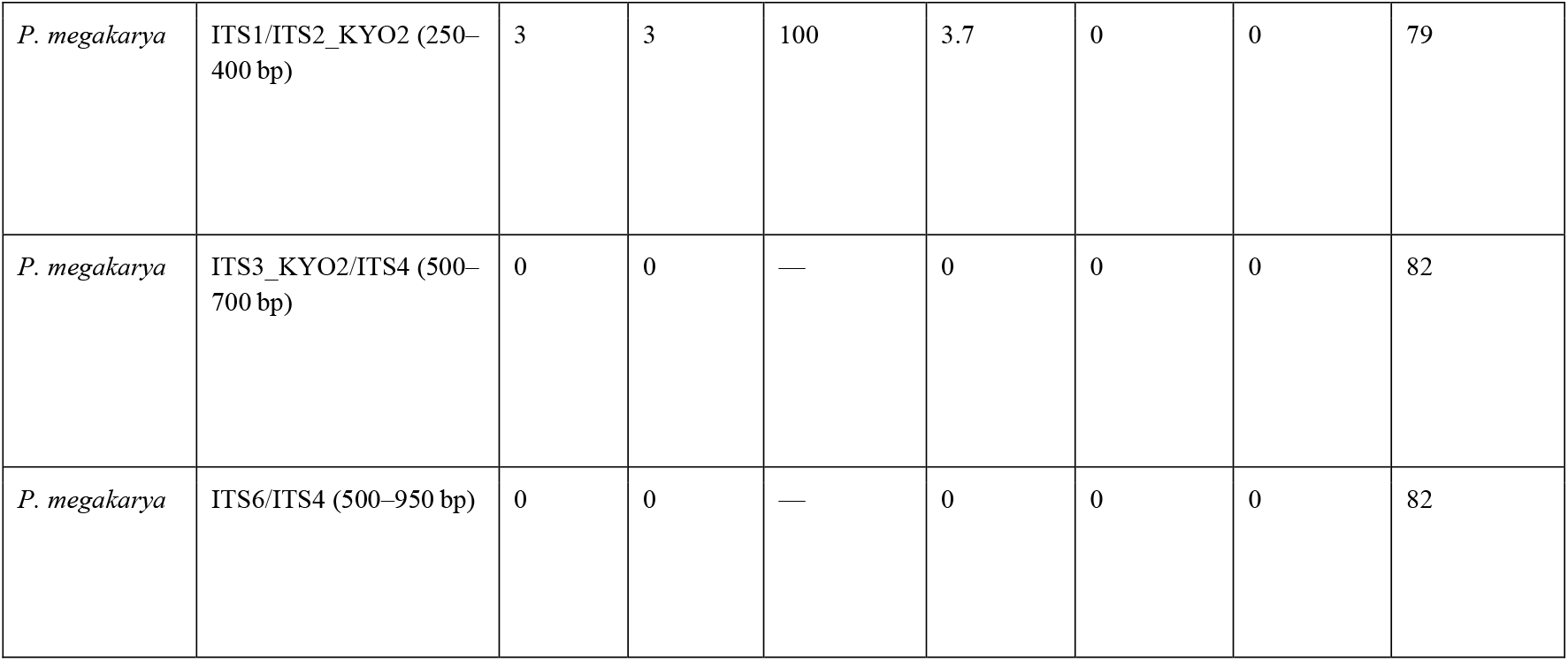
Recommended primer pairs and strict coverage by species. Coverage is reported as % of eligible under strict rules (≤ 2 total mismatches; no 3′-terminal mismatch; product within window). Ineligible counts reflect missing primer sites and/or insufficient LSU for ITS4. Complete pair matrix and ineligibility reasons appear in Supplementary Data 1. (fITS7 ≡ gITS7; Monp-ITS4R appears in the decision tree but is not tallied across all species due to limited LSU-extending *Moniliophthora* records.)

### 3.4 Mechanistic Support: Low 3′-End Entropy at Working Sites Explains Robustness

Sequence logos (information content, bits) and positional Shannon entropy clarify why the high-performing assays are stable. In *Moniliophthora*, the fITS7 forward site shows low entropy at the 3′ terminal base, consistent with strict-hit behavior whenever a compatible LSU reverse is available. In *P. palmivora*, the ITS6 forward and ITS3_KYO2 forward sites retain conserved 3′ termini; the ITS4 site is stable when present but is frequently missing, as many public records end before the LSU is reached. The entropy panels, therefore, align with the rule-based outcomes (pass/no-pass under a no-3′-mismatch constraint). Figure 2 displays the four exemplars with n indicated in each panel title; the dashed vertical line marks the 3′ base used by the rule set.

### 3.5 Rarefaction of Primer-Site Haplotypes Enables Efficient QC

Treating each binding site as a “haplotype,” rarefaction curves show how quickly primer-relevant diversity saturates. For the *P. palmivora* ITS6 site (n = 44), the mean cumulative unique count plateaus with ≈ 95% of haplotypes captured by ~ 41 sequences (50 randomizations; fixed seed = 17). Practically, ~ 40–50 accessions suffice for routine QC updates at this site; labs do not need to re-scan entire archives as new sequences accrue. We recommend re-estimating N_95_ when datasets change materially (Figure 3). A lightweight sensitivity analysis (≤3 total mismatches or allowing a single 3′ mismatch) did not alter assay rankings; details are provided in Table S3 and Figure S1.

### 3.6 A Field-Ready Decision Tree with One Cycling Program

Synthesizing coverage, entropy, and rarefaction, we recommend a two-branch decision tree operable under a single temperature program (62→45°C; 30 cycles). Temperatures are panel-specific and should be re-optimized per lab, adhering to the touchdown principle of Don et al. (Don et al., 1991). Where fungal target traits indicate the upper branch, run ITS1/ITS2 (~ 300 bp); where LSU is available or a reflex is needed, use fITS7 + Monp-ITS4R (250–350 bp). Where differential diagnosis requires excluding non-fungal agents with similar symptoms, run ITS1/ITS2_KYO2 (~ 300 bp) and optionally add ITS3_KYO2/ITS4 (550–600 bp) when LSU is present. A two-program fixed-anneal alternative (Program A 54–55 °C for ITS1; Program B 45– 48 °C for degenerate/ITS2) appears in Figure S2 and Figure 4.

## 4 Discussion

Much of what appears in the literature as “primer failure” for ITS diagnostics is an accounting problem: when public accessions truncate before the LSU region, assays that require ITS4 are often deemed ineligible, rather than being intrinsically weak, so naive tallies confound primer performance with archive incompleteness (Nilsson et al., 2008; Schoch et al., 2012). By separating eligibility (both sites present; plausible product) from performance (≤ 2 total mismatches, no 3′-terminal mismatch, size within a window), we recover a clean, mechanistic picture of what works and why. This eligibility-aware accounting avoids a recurrent denominator error in barcode benchmarking and produces a fully auditable result at the accession level (Bokulich & Mills, 2013; Taylor et al., 2016).

Under these conservative, bench-realistic rules, short ITS1 assays emerge as the most reliable defaults. For *Moniliophthora*, the ITS1/ITS2 pair covers nearly all eligible accessions. In contrast, long ITS2-forward + ITS4 assays succeed only when the LSU sequence is present, a pattern that reflects the archive structure rather than primer design (Gardes & Bruns, 1993; White, 1990). For oomycetes, ITS1/ITS2_KYO2 is highly effective among eligible *P. palmivora* accessions, while the classic ITS6/ITS4 is constrained by a very small eligible pool (Cooke et al., 2000; Toju et al., 2012). The mechanistic basis for these outcomes is the well-known inhibitory effect of 3′-terminal mismatches on polymerase extension, a principle our “no 3′-terminal mismatch” rule directly encodes (Kwok et al., 1990; Stadhouders et al., 2010). This is visually confirmed in our binding-site logos, where working assays exhibit low positional entropy at their 3′ termini (Schneider & Stephens, 1990; Tareen & Kinney, 2020).

Operationally, the recommended panel reduces complexity to a single touchdown program, a standard strategy that absorbs variations in primer melting temperature and degeneracy without proliferating protocols (Don et al., 1991; Rose et al., 1998). For quality control, treating each binding site as a “haplotype” yields a rarefaction curve that allows for principled down-sampling, meaning labs need not rescan entire archives to maintain the panel (Chao & Jost, 2012; Gotelli & Colwell, 2001).

Although demonstrated on cacao pathogens, the framework’s logic is barcode-agnostic and transferable to other systems where public records are incomplete, such as COI in animals or rbcL/matK in plants (CBOL Plant Working Group et al., 2009; Marquina et al., 2019; Paul D. N. Hebert et al., 2003). Methodologically, our principal contribution is a generalizable accounting framework rather than a system-specific primer set. Demonstrating the existence and magnitude of denominator errors under naive benchmarking, and providing auditable eligibility/performance calls, constitutes the core resource for the community. We therefore recommend minimum reporting standards for fungal ITS benchmarking: (i) eligibility and naive coverage reported side-by-side; (ii) explicit 3′-mismatch rules; and (iii) accession-level call tables and figure-generation code made public at acceptance.

Three limitations, however, merit note. First, public data for *P. megakarya* remains sparse, highlighting a need for targeted sequencing in its native range (Ali et al., 2016; Holmes et al., 2003). Second, our rule stringency is conservative, but the software allows these thresholds to be relaxed. Third, while our ‘field-ready’ panel is supported by *in silico* predictions, its efficacy should be confirmed through comprehensive bench-level validation using a diverse set of physical samples and PCR conditions before widespread field deployment. A deeper exploration of these biological and logistical variables would be a valuable next step.

In conclusion, our “eligibility-aware” framework resolves a recurrent denominator error in in silico diagnostics by separating primer performance from database artifacts, providing an accurate and reproducible template for benchmarking. This approach, combining 3’-end stringency with rarefaction-based quality control, yields a field-ready diagnostic panel for cacao pathogens. Looking ahead, our work highlights the need for new community standards. We recommend that future benchmarking papers report both eligibility-aware and naive coverages and that researchers prioritize depositing ITS sequences that extend sufficiently into the LSU region to anchor reverse primers. Adopting these transparent and auditable practices will dramatically improve the reliability of all fungal diagnostics.

## Author contributions

E.A. conceptualized and supervised the project. S.L., I.B., and M.C. developed the methodology. S.L., I.B., J.B., and E.A. performed the formal analysis. R.K.U., S.O., M.S.K., and L.M. provided resources. E.A., L.W.M., and M.S.K. acquired funding. S.L. and E.A. wrote the original manuscript draft, and all authors contributed to reviewing and editing the final version.

## Acknowledgements

Mention of any trade names or commercial products in this article is solely for the purpose of providing specific information and does not imply recommendation or endorsement by the U. S. Department of Agriculture. USDA is an equal opportunity provider and employer, and all agency services are available without discrimination. The authors used AI-based language tools (ChatGPT by OpenAI and Gemini by Google) only for grammar and clarity during manuscript editing. This work was supported by the U.S. Department of Agriculture, Agricultural Research Service, In-House Projects Nos. 8042-21220-258-000-D and 8072-41000-113-000-D.

## Data Accessibility Statement

The complete Python script used to perform all analyses is publicly available in a GitHub repository [https://github.com/EJSAHN/eligibility-aware-its-pipeline/tree/main]. All datasets generated during the current study, including accession-level call sheets, summary tables, and ineligibility breakdowns, are available as additional Supplementary Data 1 with this article. For convenience, the exact FASTA files (including GenBank accession numbers) used for the analysis are also included in Supplementary Data 2.

## Supplementary Information

**FIGURE S1.**
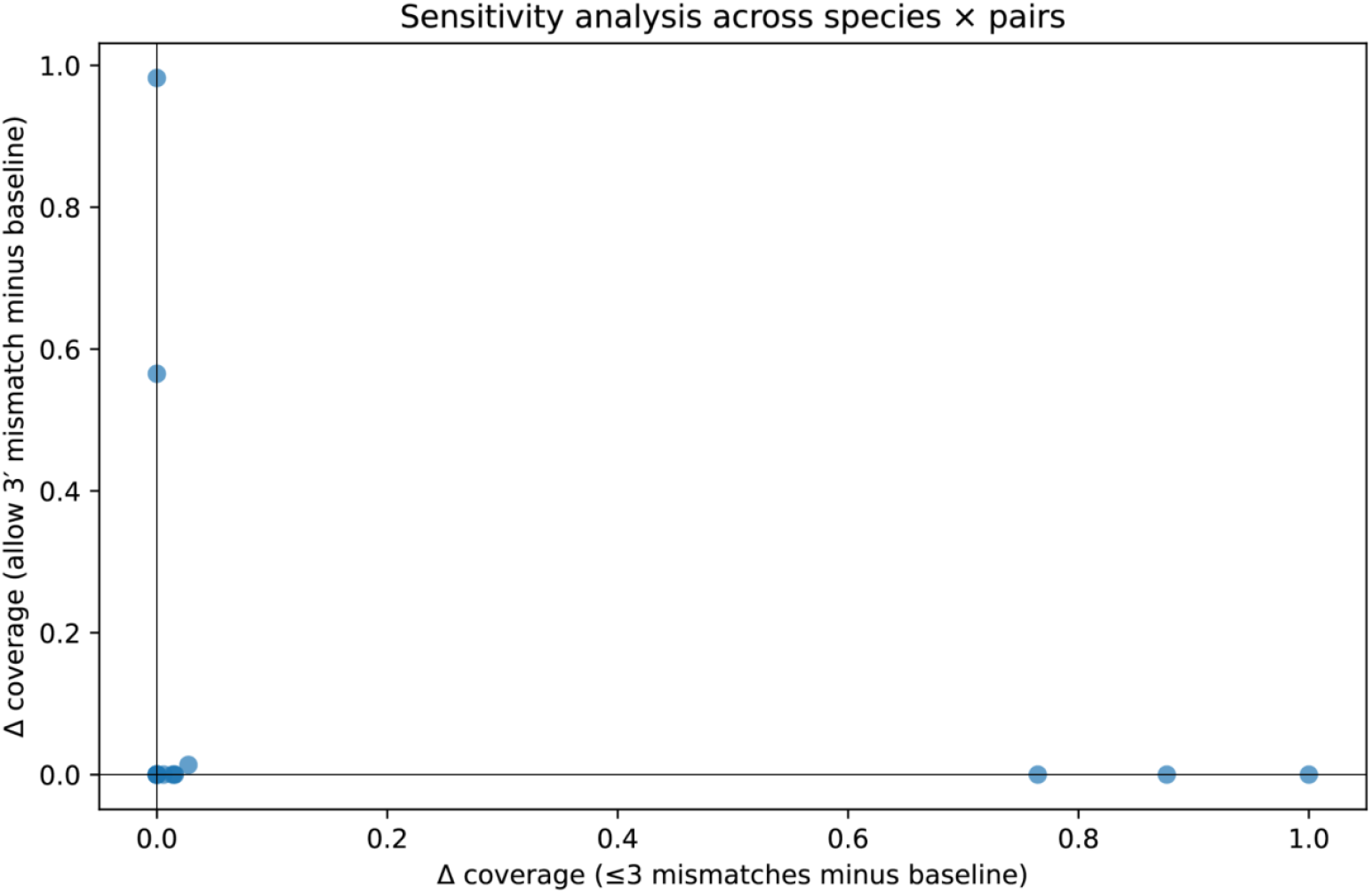
Sensitivity of eligibility-aware coverage to scoring thresholds. Left: baseline (≤ 2 total mismatches; no 3′-terminal mismatch). Middle: ≤ 3 total mismatches. Right: ≤ 2 total mismatches but allowing a single 3′ mismatch. Coverage increases modestly under ≤ 3 total mismatches without changing assay rankings; permitting a 3′ mismatch yields larger gains at a minority of sites, but relaxes a failure mode known to depress amplification at the bench.

**FIGURE S2.**
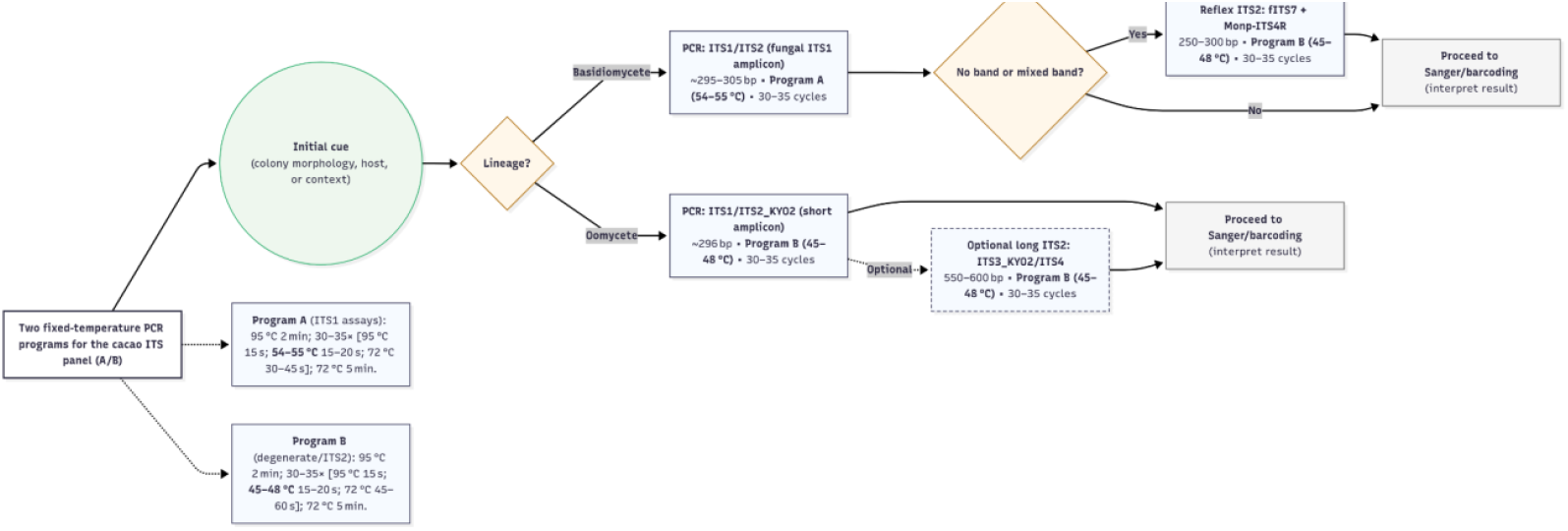
Two fixed-temperature PCR programs (A/B) for the cacao ITS panel. Program A (ITS1 assays): 95 °C 2 min; 30–35× [95 °C 15 s; 54–55 °C 15–20 s; 72 °C 30–45 s]; 72 °C 5 min. Use for ITS1/ITS2 (fungal ITS1 amplicon) and, where specificity is prioritized, ITS1/ITS2_KYO2. Program B (degenerate/ITS2 assays): 95 °C 2 min; 30–35× [95 °C 15 s; 45–48 °C 15–20 s; 72 °C 45–60 s]; 72 °C 5 min. Use for ITS3_KYO2/ITS4, fITS7/Monp-ITS4R, and fITS7/ITS4 (when LSU extends past the ITS4 site); consider Program B for ITS1/ITS2_KYO2 if Program A under-amplifies. Temperatures were chosen using the NEB Tm Calculator (OneTaq Hot Start, Standard Buffer, 200 nM).

**TABLE S1.**
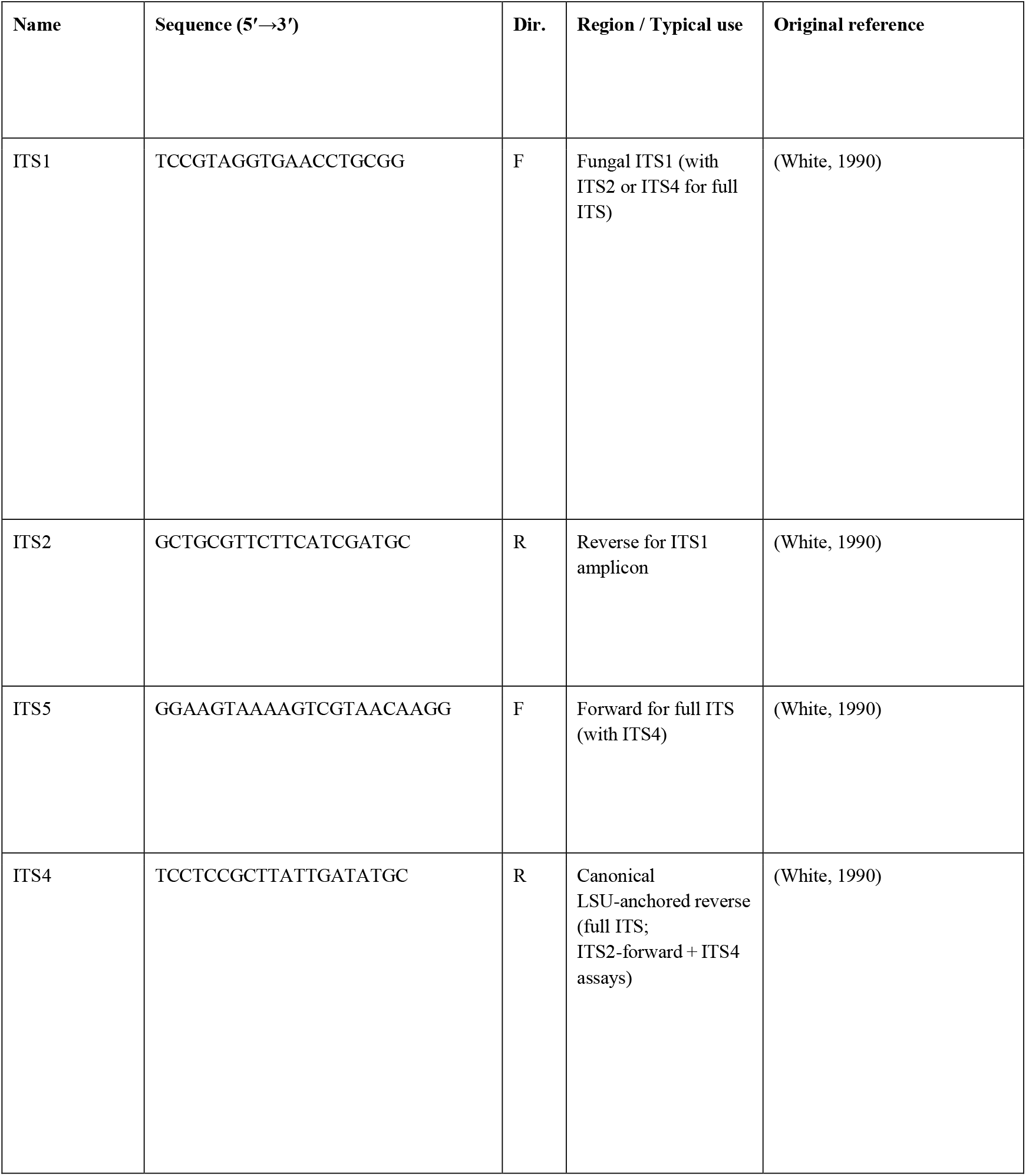
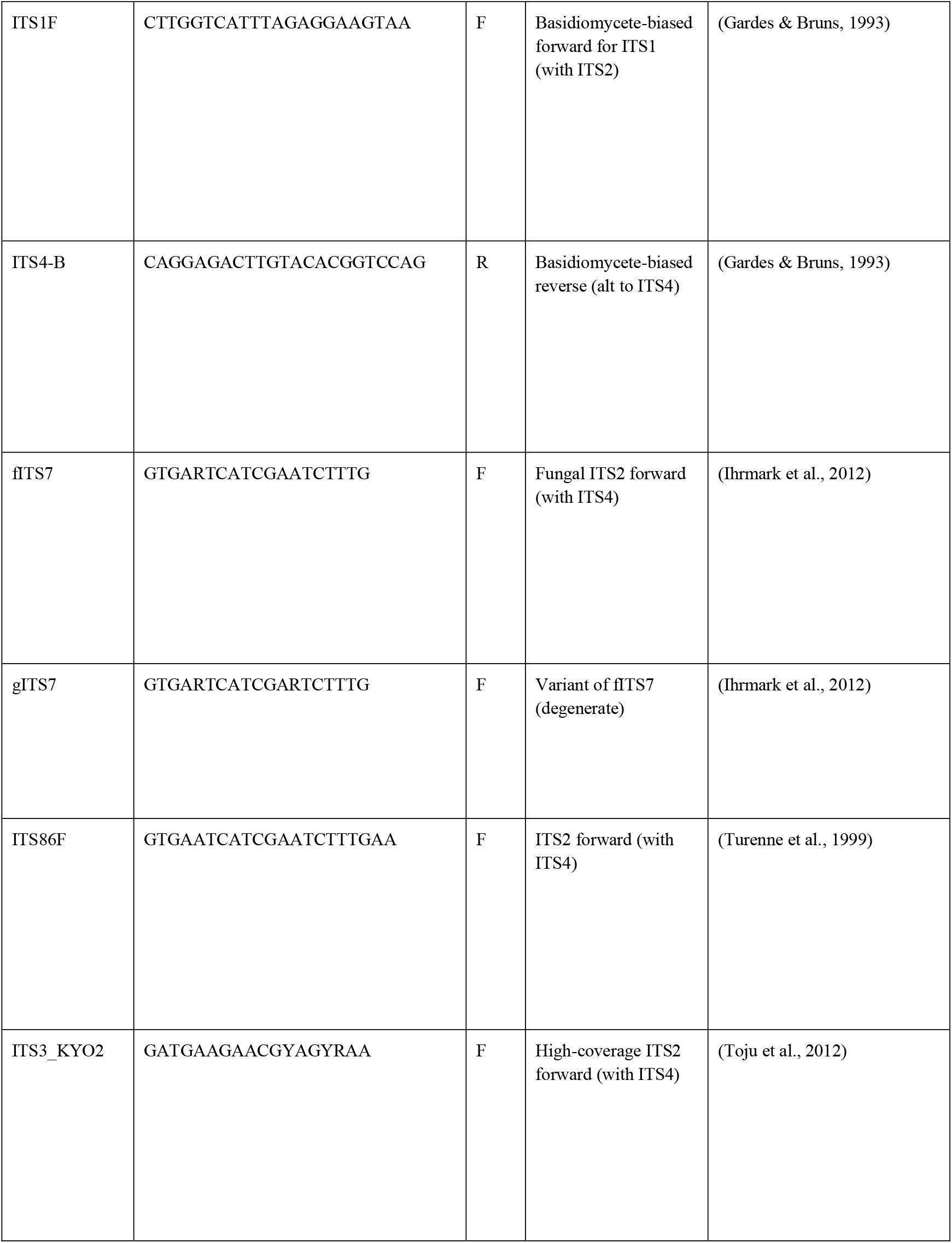
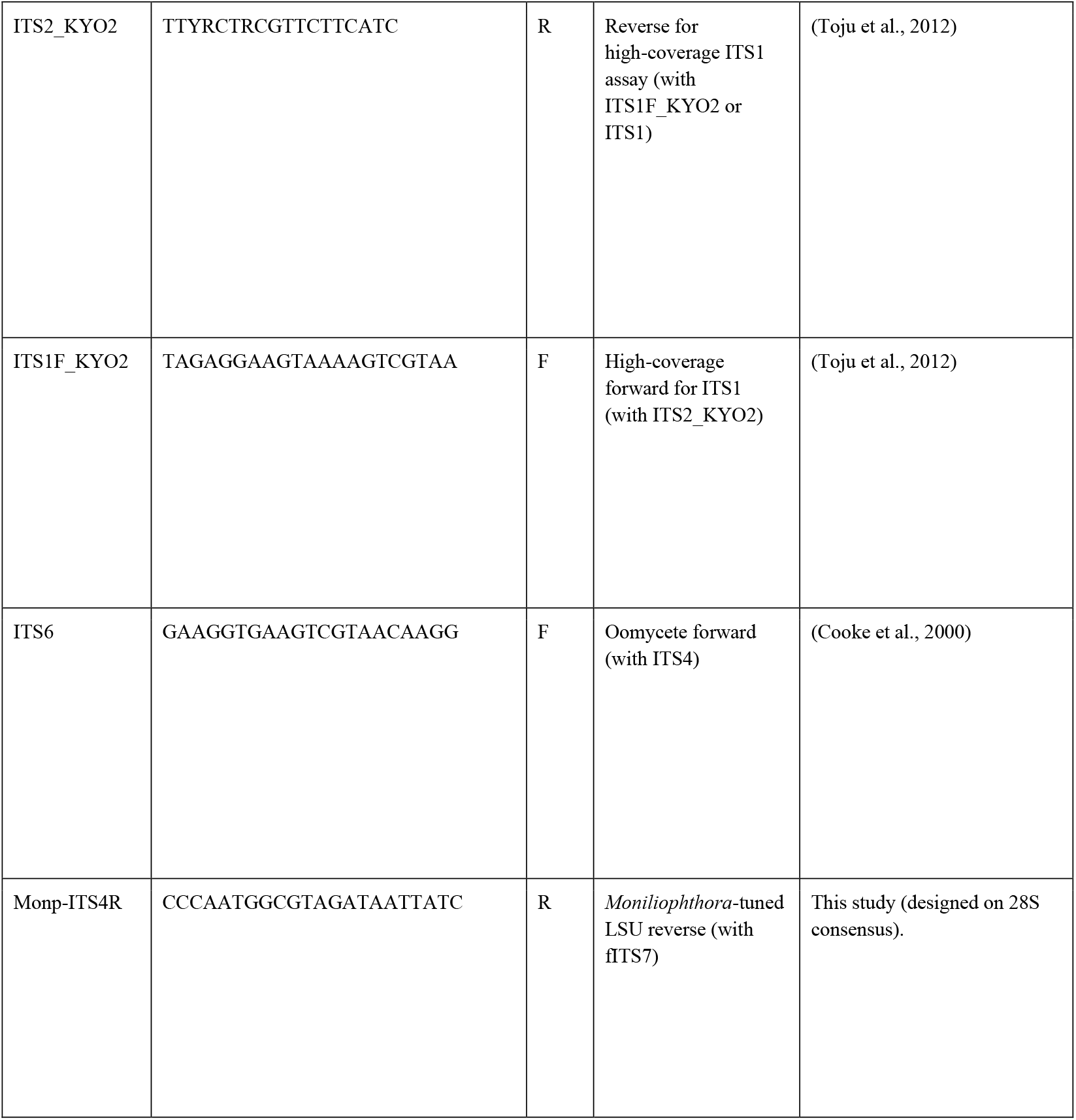
Primers used in this study. Degenerate symbols follow IUPAC conventions (R = A/G, Y = C/T, etc.). fITS7 ≡ gITS7 in usage (sequence variant). We use “ITS2-forward + ITS4” to denote forward primers that bind within ITS2 paired with ITS4; no literal “ITS2/ITS4” was evaluated because ITS2 is a reverse primer.

**TABLE S2.**
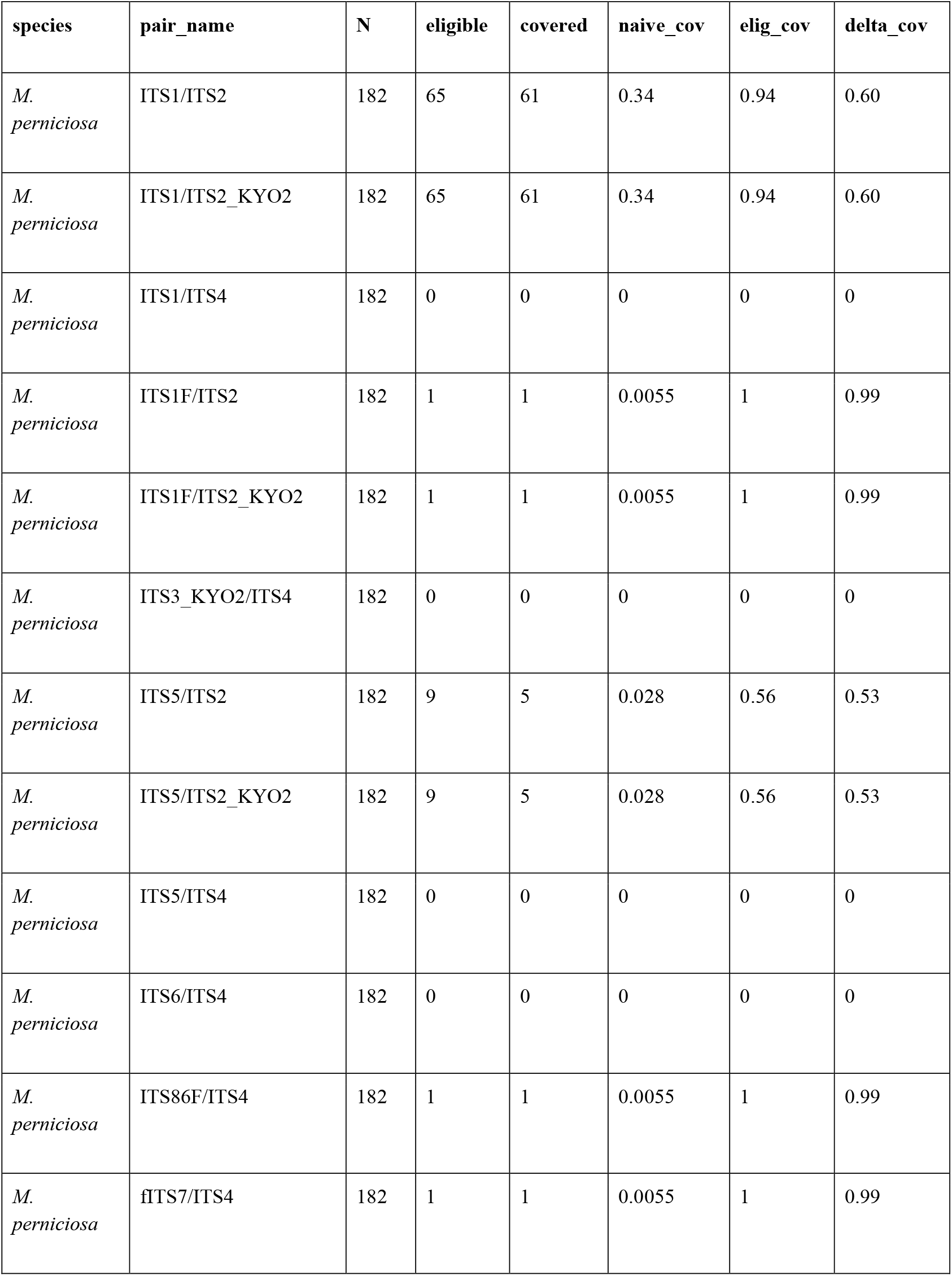
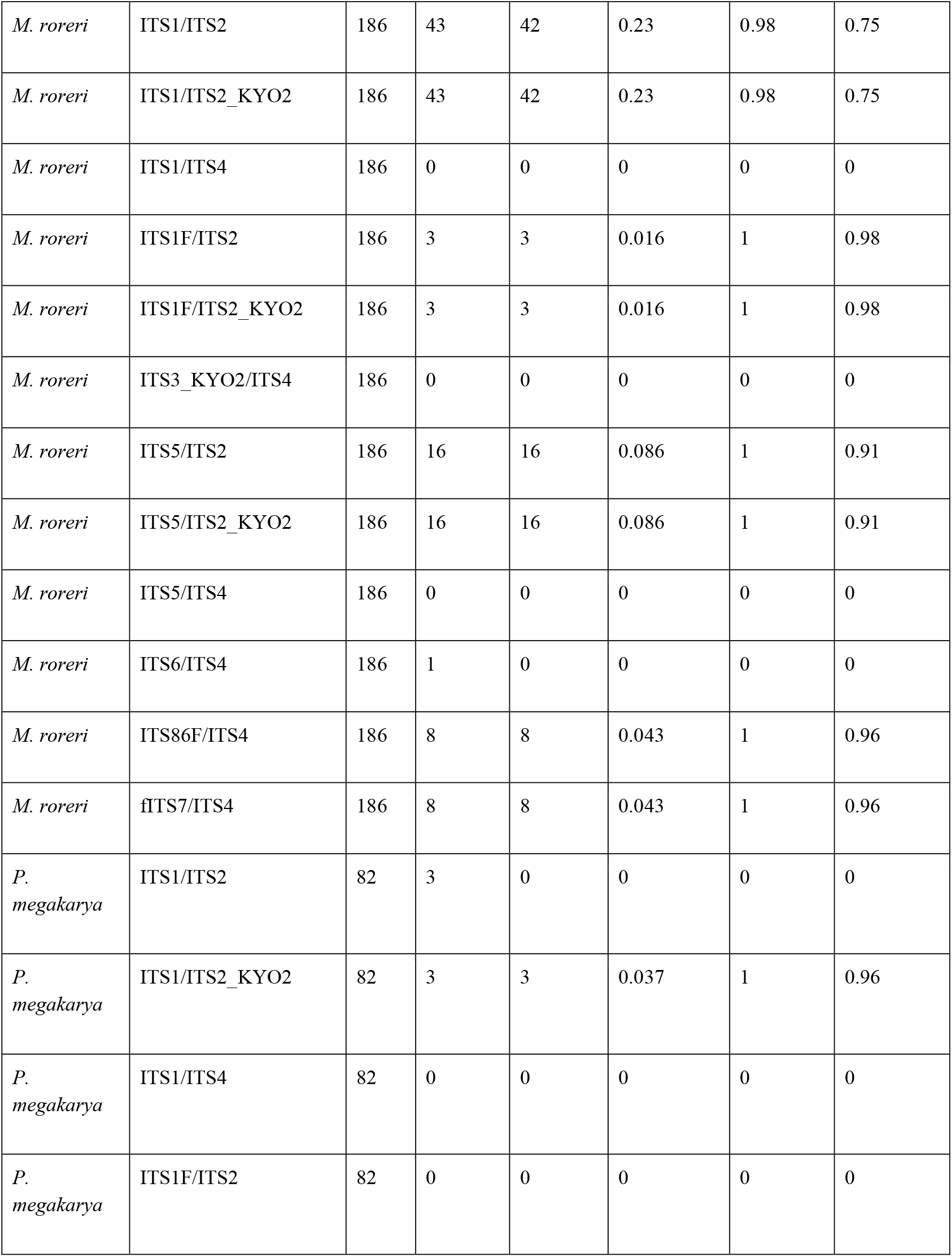
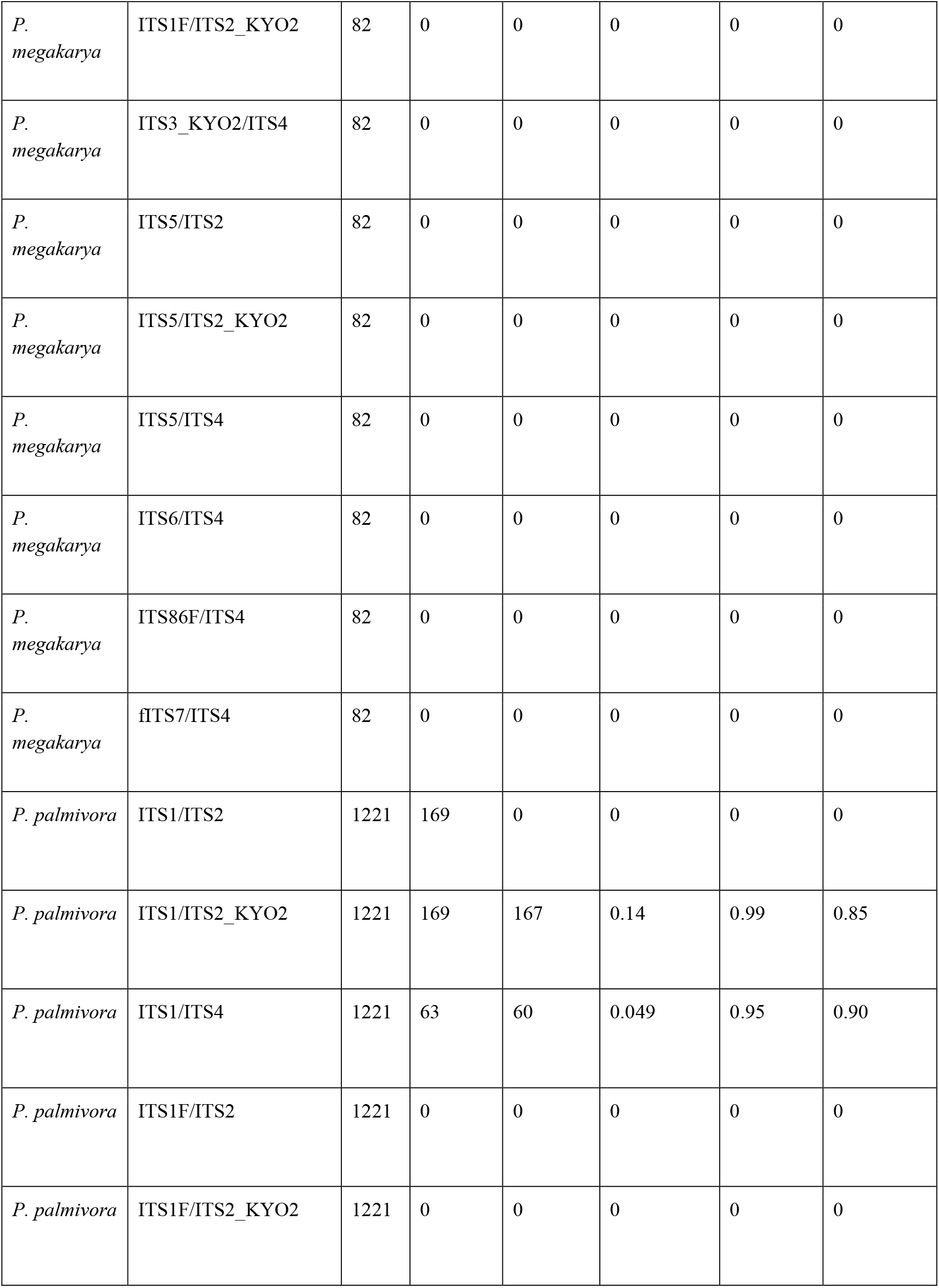
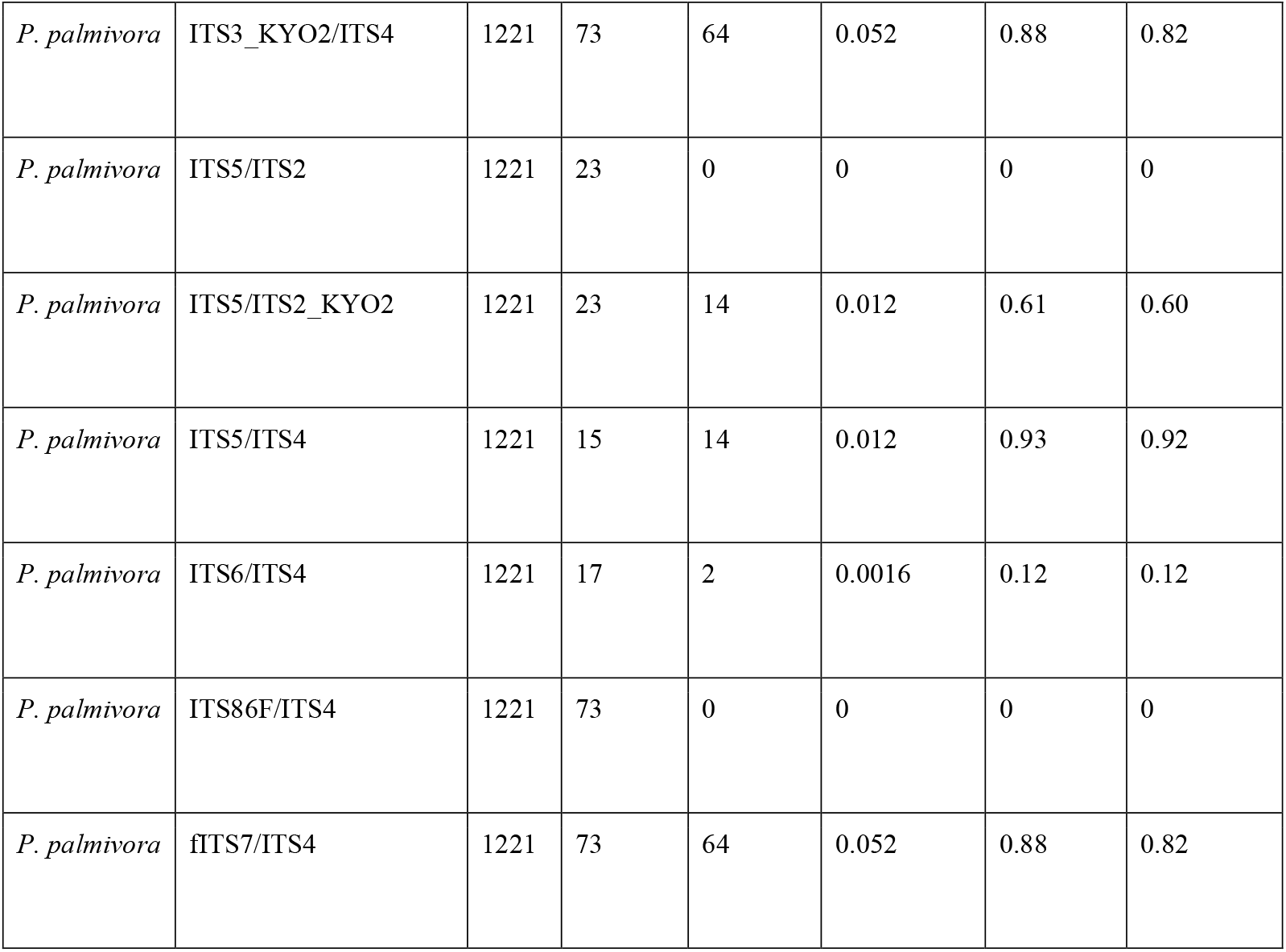
Δ-coverage: naive vs eligibility-aware coverage by species and primer pair. Δ (pp) = (covered/eligible) − (covered/N), reported in percentage points. A large positive Δ indicates that naive tallies understate primer performance because many accessions are ineligible (e.g., lack ITS4). “—” indicates undefined % of eligible when the eligible count is zero.

**TABLE S3.**
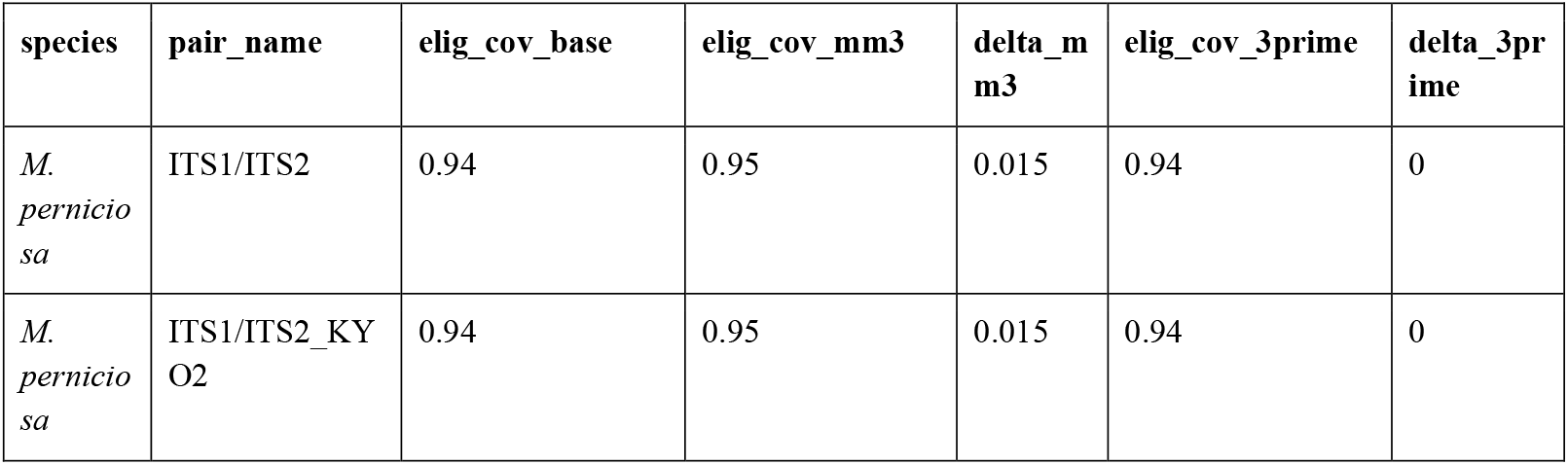
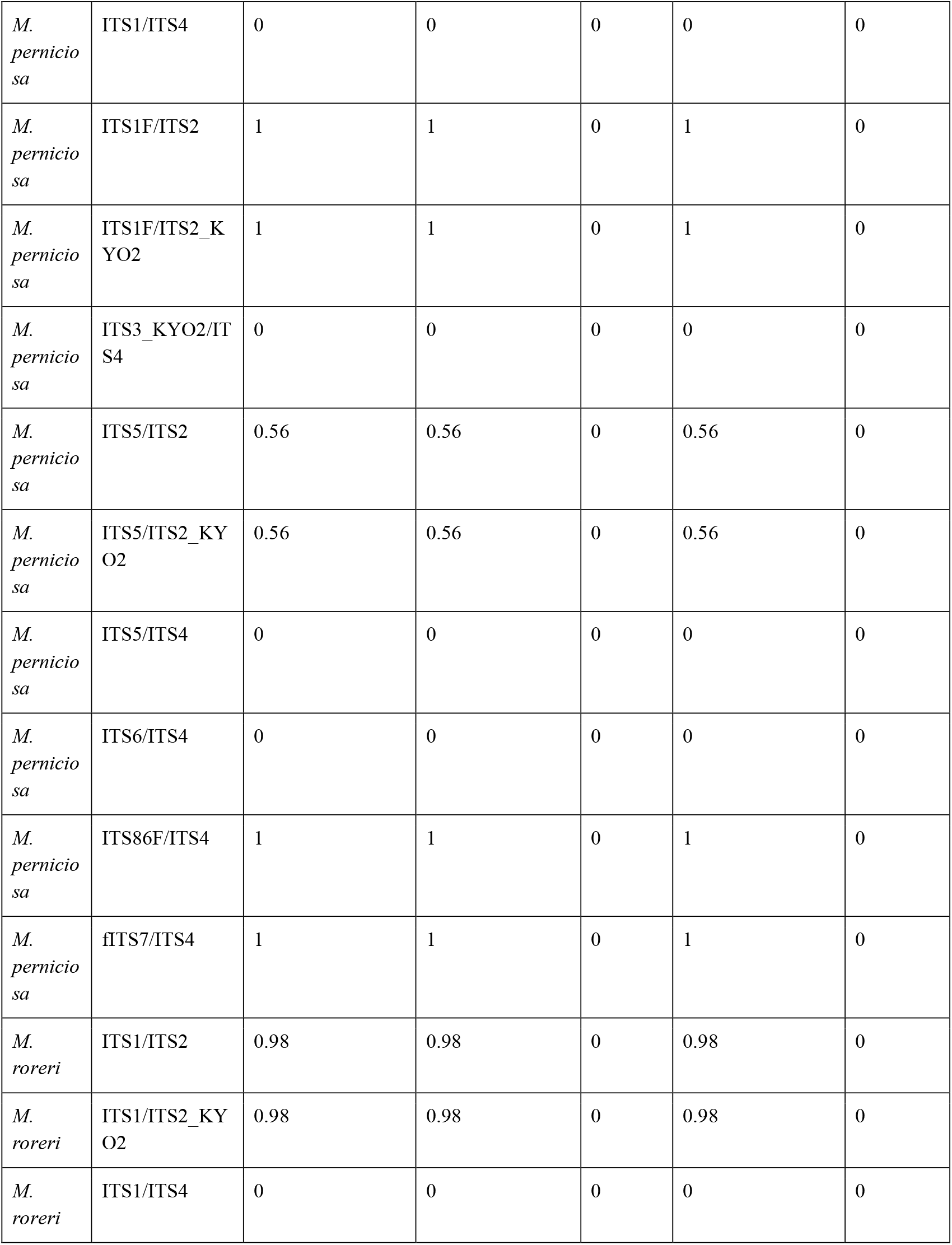
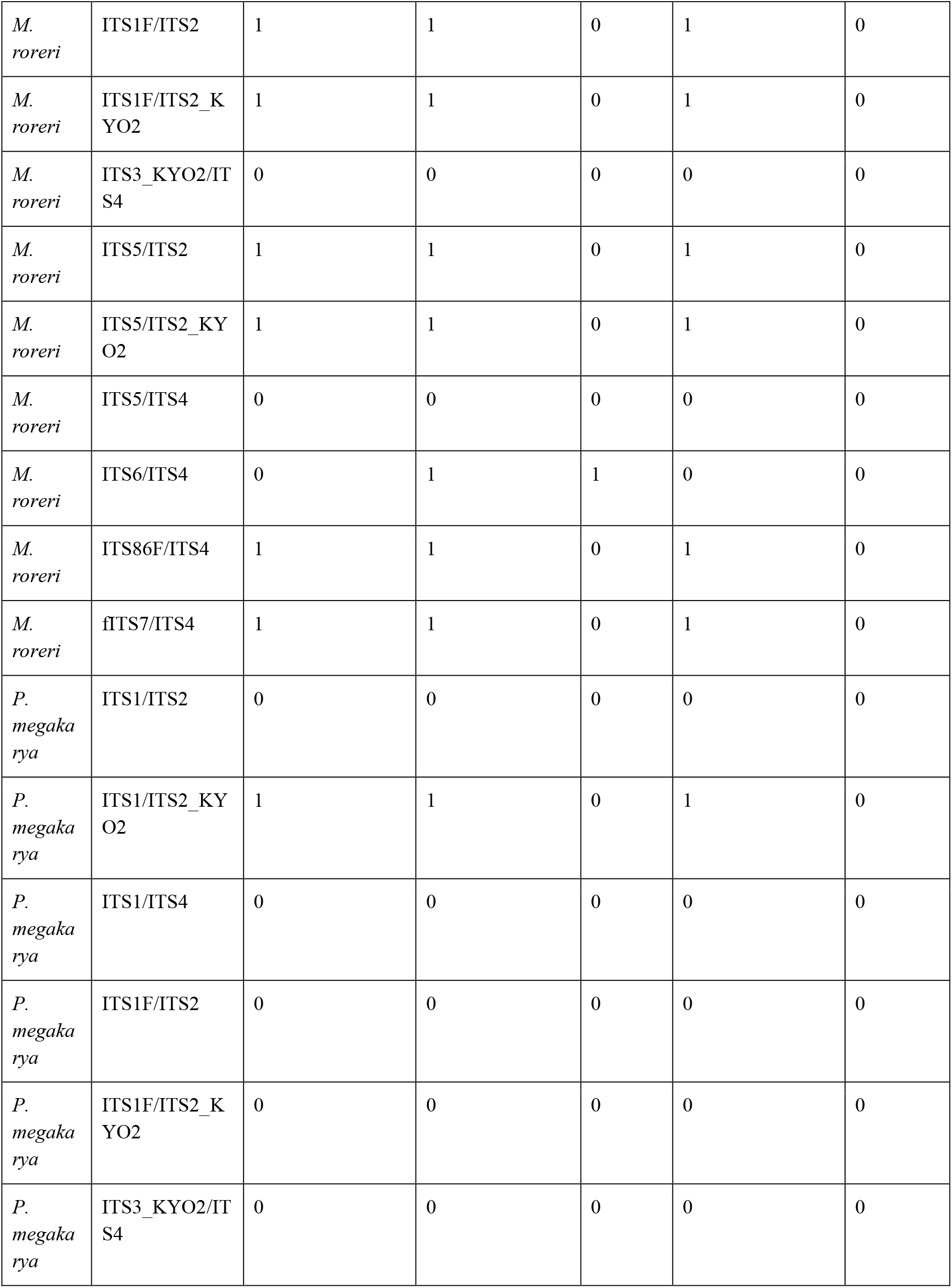
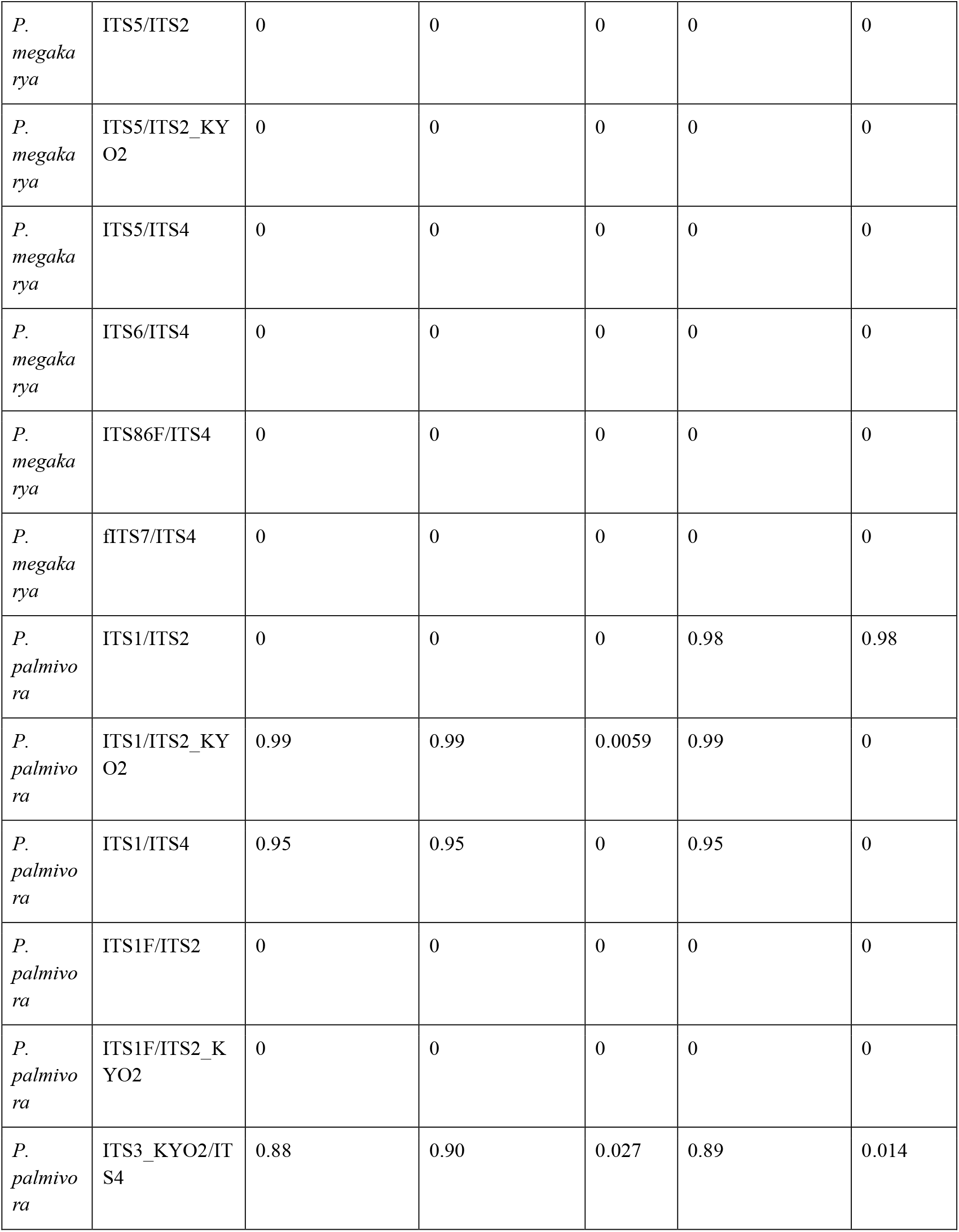
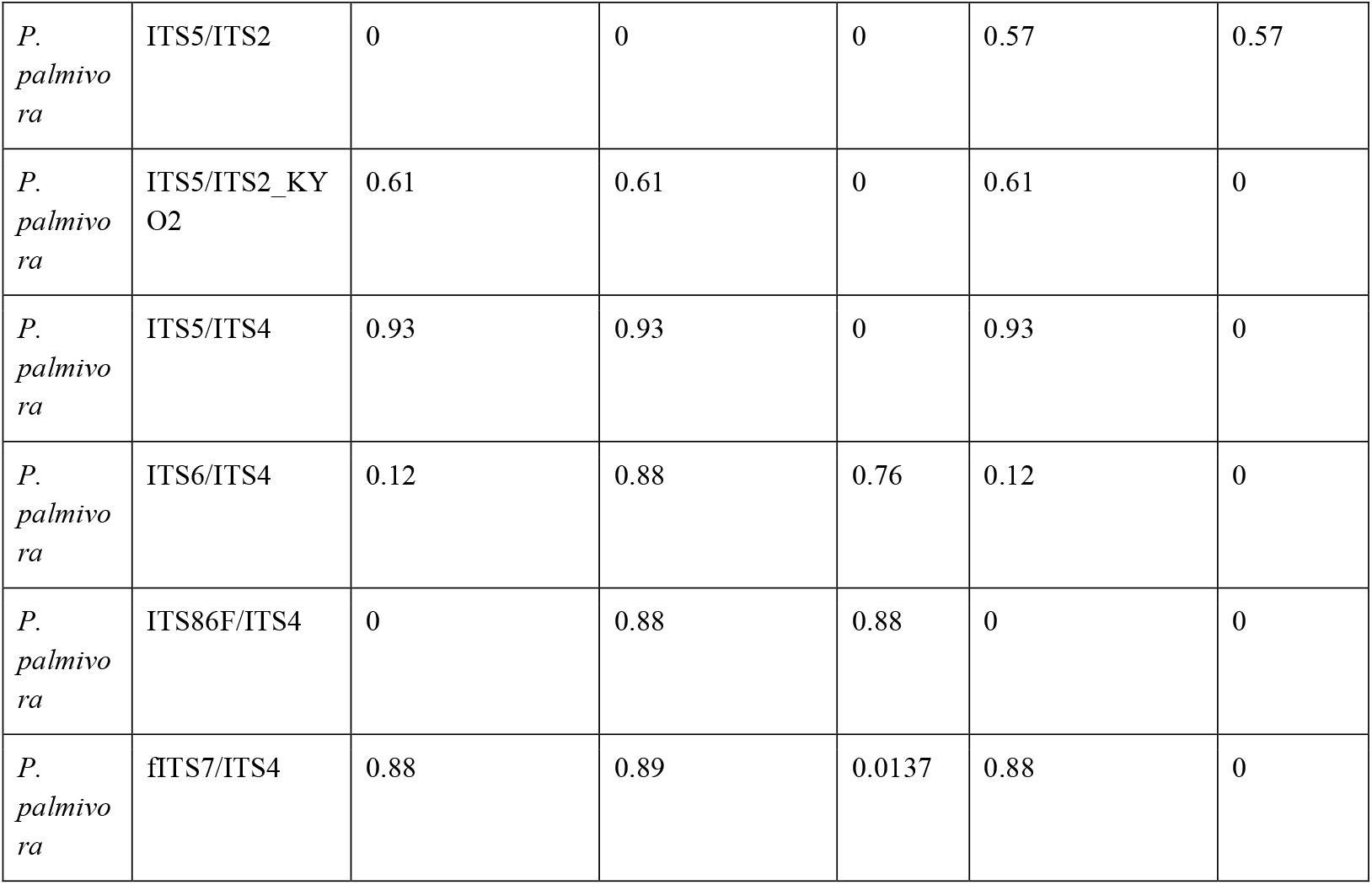
Sensitivity analysis. Eligibility-aware coverage under relaxed rules (≤ 3 total mismatches; or allowing one 3′ mismatch at ≤ 2 total mismatches). This table is provided for reproducibility and is summarized in Figure S2.

